# Phylogenetic biome niche conservatism underlies the evolution of forest paleoendemic legume trees in tropical Africa

**DOI:** 10.1101/2025.01.10.632308

**Authors:** Dario I. Ojeda, Anaïs-Pasiphae Gorel, Gilles Dauby, Sandra Cervantes, Aaron D. Pan, Thomas Marcussen, Samuel Vanden Abeele, Arthur Boom, Félix Forest, Manuel de la Estrella, Olivier J. Hardy

## Abstract

Niche conservatism is prevalent during the evolution of plant lineages. However, inferring biome niche lability and its impact on tropical tree species diversification is currently limited. To better understand biome niche lability and its effect on diversification rates, we analyzed an endemic lineage of African tropical trees, testing whether biome niche type (forest vs savanna) and biome lability are non-randomly distributed. We reconstructed a time-calibrated phylogeny of the Berlinia clade (16 genera, ca 201 species) using 140 nuclear genes, 75% of its extant species, and eight fossil calibrations. We also analyzed the phylogenetic signal of biome shifts across the group and its effect on diversification and extinction rates. We found the forest biome as the ancestral condition and a directional shift from forest to savanna (no reversals). The forest biome type is conserved in the Berlinia clade, but the lability to shift biomes is randomly distributed in the group. We did not find evidence that biome shift was associated with higher diversification or lower extinction rates. Our analyses identified five paleoendemic genera that have persisted solely in the forest biome since the Eocene. These paleoendemic and forest-restricted lineages are potentially more susceptible to habitat alterations of human related activities and future climate change, due to perhaps an innate limitation to adapt to new habitat types.

## I. Introduction

Elucidating the distribution of biome niche conservatism and niche lability (the propensity to evolve and transcend major ecological boundaries) through the evolutionary history of plant lineages can provide useful insights into their evolutionary potential under future climate change. This knowledge is particularly relevant for tropical tree lineages, which due to their longer generation times, restricted distribution and habitat specificity might render these species more susceptible to changing climatic conditions. Previous studies at regional [1,2] and global levels [3,4] suggest that biome niche conservatism has been prevalent during angiosperm evolution. The resulting phylogenetic niche conservatism (PNC) is reflected in the consistency of the climate niche of tropical forest tree lineages along similar rainfall gradients across continents [5]. These studies indicate that PNC have played a major role shaping the current distribution of plant diversity.

Biomes capture major vegetation types defined by climate, life form and ecophysiology. Shifts among biome types are considered major transitions involving important adaptations [6]. Understanding these transitions are particularly relevant in regions where major changes in biome types have occurred in the past over wide evolutionary time frames, under predicted human disturbances, and future climate change [7]. In Africa, the tropical rainforest biome has experienced an overall decline of area since its maximum extension in the early Eocene to the Middle Miocene (55-11 Ma) [8]. The contraction of the forest biome has been proposed as one of the reasons for the depauperate species diversity of many tropical tree lineages in Africa (e.g., palms and some legume lineages) [9] compared to their counterparts in Asia and the Neotropical regions [10,11]. On the other hand, the overall decrease of rainforest area since the Eocene in tropical Africa has also made new habitat types available (e.g., savanna biome), created opportunities for biome shifts, and changes in diversification rates. In spite of this, a recent analysis suggested that niche conservatism of biomes (forest and savanna) is prevalent in most (93%) speciation events in tropical African tree species at the genus level [12]. Indeed, evidence shows that several arborescent forest taxa contracted and expanded their distribution range with the cycles of rainforest changes (without shifting biomes), following the extension of the forest biome [5,13,14]. Thus, despite these repeated forest biome contraction/expansion cycles and the availability of new biomes, most African tree lineages have maintained their biome across wide evolutionary time frames. It thus seems that the capacity to shift between the forest and savanna biomes is unevenly distributed across clades [12]. This suggests that some lineages could be more susceptible to higher extinction rates when the availability of these biomes is altered. However, understanding how biome niche lability is distributed among lineages and the timing of their shifts require robust time-calibrated phylogenetic reconstructions at the species level with good representation of extant taxa. We currently lack this knowledge for most tropical trees at the local and global levels, particularly for tree lineages that are endemic to the African tropical region.

In Africa, the Guineo-Congolian lowland rainforest has been identified as a region of plant paleoendemism [15,16]. Paleoendemic genera (taxa that diverged early and with a restricted distribution) have been previously suggested in the Berlinia clade (Subfamily Detarioideae, Leguminosae) [17], a group of tropical tree legumes that diversified in the Guineo-Congolia region. The Berlinia clade consists of 16 genera, nine of which are composed of at least 10 species, totaling ca. 201 species [18]. Unlike other African forest tree taxa, the Berlinia clade seems to have experienced an increase in diversification rates since the Miocene [17]. The most recent analyses within this clade indicate different diversification patterns, where some lineages seem to have exploited niches within the forest biome, while others were able to exploit the new biomes, such as open grasslands and woodlands, that became available as the forest cover area reduced [17,19]. However, limited representation of the group has hindered the reconstruction of the distribution of biome niche lability and whether this capacity has led to higher diversification and/or extinction rates.

To understand more in depth the distribution of biome niche conservatism and lability, we performed a phylogenomic reconstruction of all 16 genera of the Berlinia clade, including 75% of all extant species. We generated a time-calibrated phylogeny using eight fossils from the Eocene onwards to obtain the phylogenetic placement and the timing of biome shifts.

Specifically, we aim to test the following hypotheses: 1) biome type shows phylogenetic signal suggesting that some lineages (genera) within the Berlinia clade have remained in their ancestral biomes despite the availability of new habitats during their evolutionary history, 2) the capacity to shift biomes (biome lability) from forest to savanna is non-randomly distributed within the Berlinia clade at the genus level, 3) lineages with the capacity to shift biomes show higher diversification rates compared to lineages that retain the ancestral forest biome, and 4) lineages that retained their ancestral forest biome have higher extinction rates compared to lineages that shifted to the savanna biome.

## 2. Materials and Methods

### (a) The study system

The Berlinia clade comprises an estimated 201 species grouped in 16 genera [20]. The entire clade is endemic to Africa and here we included 152 species, representing all 16 genera, and 75% of the species (Table S1). We used *Paramacrolobium coeruleum* as an outgroup species in all the analyses.

### (b) DNA extraction and target enrichment

DNA was extracted from leaves (25-35 mg) obtained from herbarium specimens or silica gel dried samples using a modified CTAB protocol [21] and cleaned using the QIAquick PCR Purification Kit (Qiagen, Venlo, Netherlands). We employed a modification of the protocol for plastome capture [22] for library preparation. Hybrid enrichment was performed on pools of 48 samples per reaction according to the MYbaits v2.3.1 protocol, using 23 h of hybridization, a high stringency post-hybridization wash, and a final amplification using 15 PCR cycles. The Detarioideae v.1 bait [19] was used for this study. This bait set consists of 6,565 probes (120 bp long overlapping baits) targeting 1021 exons from 289 genes. Paired-end sequencing (2 × 150 bp) was performed on an Illumina NextSeq with reagent kit V2 at the GIGA platform (Liège, Belgium), assigning ca. 400,000 reads/sample. Orthologs were recovered with the strategy developed previously [23]. First, we assembled *de novo* reads for each species using SPAdes ver. 3.9 [24] and reduced redundancy of the clusters recovered using CD-HIT [25] (99%, threshold, word size = 10). Then we performed an all-by-all blast on all the samples and later filtered with a hit fraction cut-off of 0.5. We applied MCL [26] using an inflation value of 1.4 to reduce identified clusters in the samples. Clusters with less than 1,000 sequences were aligned with MAFFT [27] (-genafpair-maxiterate 1,000, and 0.1 minimal column occupancy) and tree inference was generated with RAxML v. 8.2.9 [28]. For larger clusters (> 100 sequences) we used PASTA [29] with minimal column occupancy of 0.01 and trees inferred using fasttree [30]. Finally, orthologues were selected using the strict one-to-one strategy. This final step allows the identification and exclusion of paralog sequences, which is an advantage over other pipelines that only identify these paralog regions [31,32]. The matrix was inspected and formatted with AliView ver. 1.28 [33].

### (c) Phylogenomic reconstruction with individual orthologs, concatenated and multispecies coalescent approaches

We performed phylogenetic analyses on each separate ortholog (gene tree) and on the concatenated matrix (supermatrix) with maximum likelihood (ML) as implemented in RAxML ver. 8.2.9 [28] using the GTRCAT model with-f a flags, 1,000 bootstraps, and default settings. We also carried out a ML analysis as implemented in IQ-TREE 1.6.1 [34] with ultrafast likelihood bootstrap with 1,000 replicates. In addition, we carried out a Bayesian analysis using MrBayes 3.2.6 [35,36] on the concatenated matrix. For the Bayesian analyses we applied four chains, two runs of 50,000,000 generations with the invgamma rate of variation, and a sample frequency of 5,000. The performance of the Bayesian analysis was assessed with Tracer 1.7 [37]. The species tree estimation was performed under the coalescent model using the individual ML gene trees obtained with RAxML to infer a species tree with ASTRAL-II v. 5.5.7 [38]. Support was estimated using local posterior probability (LPP). We used *Paramacrolobium coeruleum* as outgroup taxon in all inferences. The obtained trees from all the analyses were visualized and edited with FigTree (Rambaut, 2016).

### (d) Climate niche modeling and categorization of biomes

To categorize biomes within the Berlinia clade, we used a database of 4,142 tree species across tropical Africa extracted from the RAINBIO project [39]. The bioclimatic group of the 152 sequenced species from the Berlinia clade were inferred from georeferenced coordinates and predicted using niche similarity, a trait that encompasses the wide range of strategies allowing plants to persist in specific climatic conditions [12]. This modeling resulted in the classification of two biomes, which corresponded to the forest and savanna biomes. The biome assignment of the Berlinia species was further corroborated with literature, descriptions from herbarium specimens, and from our own field expertise of the distribution of the species. Biomes were further corroborated with recent taxonomic revisions for all representatives in *Anthonotha* [40], *Englerodendron* [41], *Microberlinia* [42], and the monotypic genera *Icuria* [43], *Michelsonia* [42], and *Librevillea* [44,45] (Table S3).

### (e) Ancestral state reconstruction and biome shifts

We mapped the two biomes at the species level to reconstruct ancestral states within the Berlinia clade and to infer the number of biome shifts. Biomes and ancestral state reconstructions were analyzed on the best RAxML tree recovered from the concatenated dataset. These reconstructions were performed using parsimony and likelihood methods as implemented in Mesquite ver. 2.75 [46]. The number of inferred biomes shifts from this analysis was later used to estimate the level of aggregation of biome lability (see below). We estimated the delta statistic (δ) as a measure of phylogenetic signal [47] for biome (forest and savanna) using the “delta” function in the R package ‘ape’ version 5.3 [48]. Delta measures the degree of phylogenetic signal between any given categorical trait and a phylogeny of the group of interest. The obtained δ-value was then compared to the δ-values obtained from 100 random permutations of the tips in our phylogenetic tree. A p-value less than the level of the test (0.05) means phylogenetic signal between the trait and the tree. We used this p-value to determine if biome type has a phylogenetic signal in this group. In addition, we assessed the phylogenetic signal by estimating D, a metric based on the sum of sister-clade differences in the phylogeny [49], using the R package caper. We performed 1000 permutations and compared the probability under a random (D = 1) phylogenetic structure. Finally, we evaluated the level of aggregation of biome lability (labile vs. non labile) using the mean phylogenetic distance between the number of instances where this trait evolved in the Berlinia clade using the tree we obtained from the BEAST calibration. First, we first converted to an ultrametric tree using the chronos function in ape (lambda = 0). We tested the null hypothesis that the instances of biome lability at the genus level (six nodes for the genera *Julbernardia, Bikinia, Aphanocalyx, Brachystegia, Berlinia*, and *Isoberlinia*) in the Berlinia clade were aggregated compared to 1000 random set of nodes using a custom-made R script.

### (f) Divergence time estimates

There are several fossils assigned to extant genera of the Berlinia clade and we employed a total of eight fossils to constrain the ages of internal nodes within the clade (Fig. 1 and Table S5) [50–57]. Divergence times were estimated with BEAST v. 1.10.3 [58] under a HYK substitution model with strict clock, coalescence constant-size, and a speciation Yule process, and a lognormal distribution for the calibration points. We employed eight fossils to generate priors of stem node age for four nodes in the topology (in subclades A and B, Table S5). These priors were generated using the age of the oldest fossil and the number of fossil sites that corresponded to each lineage. Here we employed a modification within the Berlinia clade (Table S6) of the approach previously used to calculate these priors [59]. In subclade A, we used the following parameters for the *Brachystegia* stem: mu = 1.9673, sigma = 0.3993, offset = 23.03, and for the *Aphanocalyx* fossil: mu = 5.0252, sigma = 1.0939, offset = 46. In subclade B, for *Anthonotha* and *Englerodendron* fossils, we applied mu = 4.2752, sigma = 1.0939, and offset = 21.73. We ran two independent runs of 200 million generations, sampling trees and parameters every 10,000 generations, and a final burn-in of 20,000 million generations. Tracer 1.7 [37] was used to assess convergence among the chains as well as to evaluate the ESS parameter (ESS > 200).

**Figure 1.**
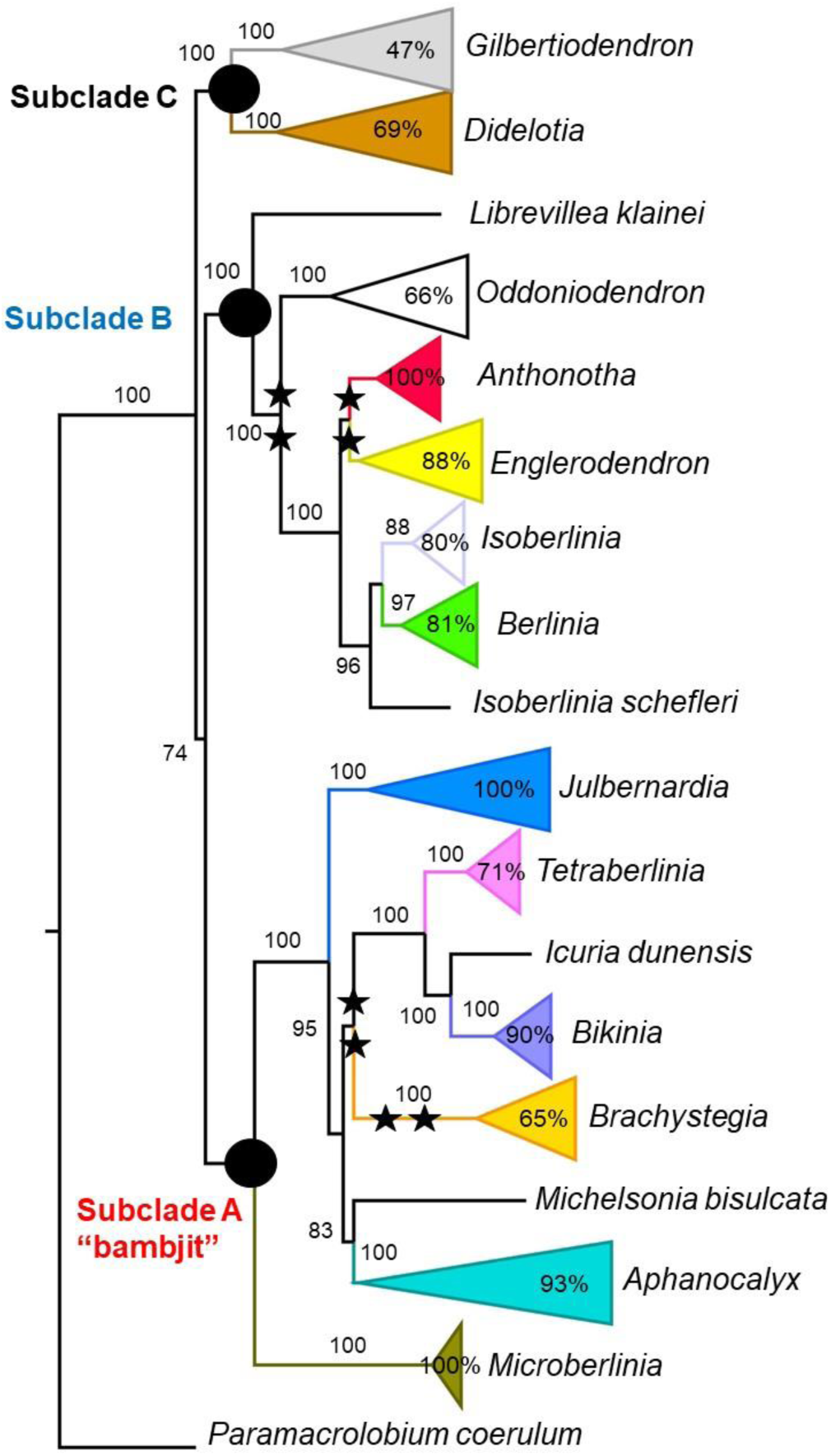
Phylogenetic relationships within the Berlinia clade based on maximum likelihood as implemented in RAxML and using the concatenated data set. The major subclades recovered within the group are marked with black circles. Percentage values of each genus represent the portion of species included. Values next to the branches represent bootstrap values. The stars indicate the locations of the eight fossil calibration points.

### (g) Diversification rates in the forest and savanna lineages

#### (i) Time-dependent diversification

To assess the variation on diversification over time, we performed a time-dependent diversification analysis as implemented in RPANDA [60]. Our dated tree covered all extant genera and 76% of the species (fraction, f = 152/201). We tested six models: 1) constant speciation rate (λcon) with no extinction (Yule null model), 2) exponential variation of speciation rate (λexp) with no extinction, 3) linear variation of speciation rate (λlin) with no extinction, 4) constant speciation and extinction rates, 5) exponential speciation rates with a constant extinction rate, and 6) linear speciation rates with a constant extinction rate [60]. These models estimate current speciation and extinction rates with their corresponding parameters by measuring their variation over time up to the crown age. The diversification rates and major shifts were also evaluated using a Bayesian approach with the reversible MCMC method as implemented in BAMM v.2.5.0 [61,62]. Our interest was to identify lineages (at the genus level) exhibiting significant rate shifts in speciation, extinction, and net diversification, and to conduct a rate-through-time analysis of these shifts within the entire Berlinia clade. Incomplete sampling of the remaining species was accounted for by specifying genus-specific sampling fractions. Priors for BAMM were generated with BAMMtools v.2.1 [61] in the R statistical environment [63]. The BAMM analysis was run with four Markov chains for 50,000,000 generations sampling every 5,000 generations. Considering the size of the phylogeny, we set the expected number of shifts to 1. The convergence was assessed by the ESS of likelihood and the number of shifts (ESS > 200). We removed 20% of the samples from the posterior distribution sampled by BAMM and analyzed the output with BAMMtools. We computed 95% credible rate shift configurations, estimated the clade (focusing on the genus level) specific evolutionary rates by obtaining rates-through-time (RTT) plots (λ, μ, and r), and obtained the visualization of the mean phylorate plots. In addition, we used the R package phytools ver. 1.5-1 [64] to generate lineage-through-time (LTT) plots to determine the temporal pattern of lineage accumulation in the Berlinia clade and its 16 genera. All tests were performed on the dated phylogeny obtained with BEAST.

#### (ii) Biome and trait-dependent diversification rates

We assessed the effect of biome shifts as potential drivers of the diversification in the Berlinia clade. We used the two biomes (forest and savanna) as previously identified in the climate niche modeling. In addition, we also explored the effect of biome lability on the diversification rates. To investigate whether biome or biome lability have promoted diversification rates in the Berlinia clade, we used the BiSSE model [65,66] where extinction and speciation rates are associated with phenotypic evolution of a trait along a phylogeny. We used the dated tree obtained with BEAST and ran four independent MCMC chains for 10 x 10^4^ steps for each of 10 randomly chosen trees using an exponential prior for the rates. We discarded the first 25% of steps of each chain as burn-in. We also tested differences in speciation rates between the two biomes and labile vs not labile using maximum likelihood (ML) and tested for significant differences among the models with ANOVA tests. Both ML and MCMC were run as implemented in the R package diversitree [66].

## 3. Results

### (a) Phylogenomic reconstructions and major lineages within the Berlinia clade

The aligned matrix of 140 nuclear DNA sequences (orthologs) consisted of 56,605 bp, of which 12,890 were variable sites and 11% were parsimony informative sites (Table S2). We recovered similar topologies using ML, Bayesian, and species tree with ASTRAL, obtaining highly resolved relationships among genera and within each of the 16 genera of the Berlinia clade (Fig. 1, Figs. S1-S5). We recovered three major lineages (subclades A-C), congruent with previous results on their taxonomic relationships [42] and with previous phylogenetic analyses based on a more limited representation of the group [17,19,41]. All genera within the Berlinia clade were recovered as monophyletic, except *Isoberlinia*, where *I. scheffleri* was found as sister to a clade comprising genus *Berlinia* sister to the remaining species of *Isoberlinia* included in this study (Fig. 1 and Figs. S1-S5).

### (b) Biome shifts within the Berlinia clade

Using a comprehensive database of over 4,000 African tree species, along with the coordinate locations of the Berlinia clade specimens used in this study, we identified two biomes corresponding to the forest and savanna. Nearly 80% of the species within the Berlinia clade were classified as inhabiting the forest biome (Table S3). We inferred the forest biome as the ancestral habitat for the Berlinia clade, as well as all its 16 genera. Biome lability was observed in six genera (*Berlinia, Julbernardia, Bikinia, Brachystegia, Isoberlinia,* and *Aphanocalyx*), with nine independent transitions from the forest to savanna, all localized in subclades A and B (Fig. 2). No reversals from the savanna to forest biome were observed. We identified two distinct patterns of biome shifts. Firstly, one shift to savanna with subsequent diversification within the savanna lineages in *Brachystegia*, *Julbernardia* and *Isoberlinia* (excluding *I. scheffleri*).

**Figure 2.**
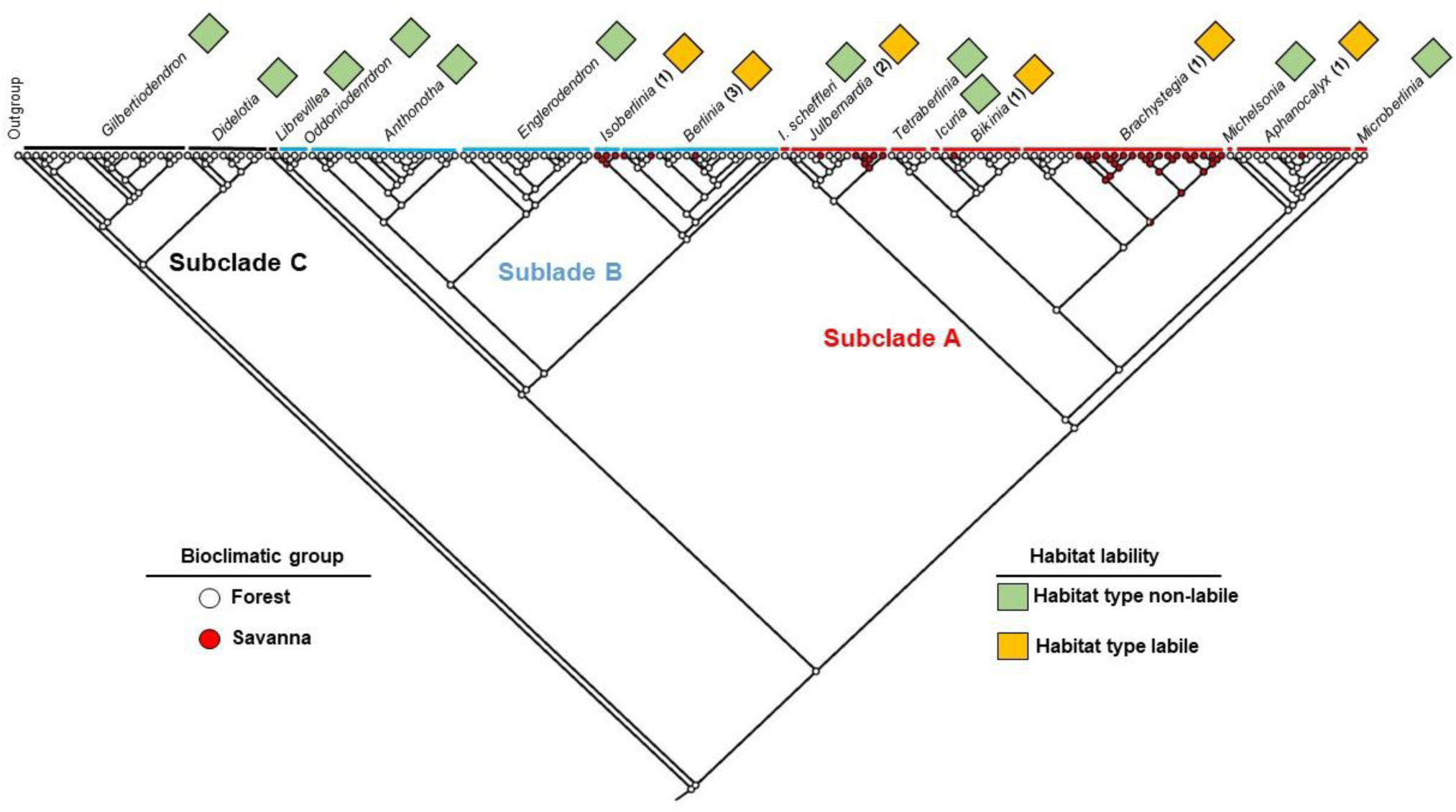
Distribution of the two bioclimatic groups and biome lability within the Berlinia clade at the genus level. The color of tips and nodes indicate the (ancestral) bioclimatic group. Green squares represent genera that remained in the forest biome, orange squares represent genera where shifts to the savanna biome occurred. The number in parenthesis indicates the number of transitions within each genus.

Secondly, one to three independent shifts to savanna within each genus, but without further diversification in the savanna lineages in *Bikinia, Aphanocalyx*, *Julbernardia*, and *Berlinia*. In *Julbernardia* we found the two previous patterns occurring at different time scales (Fig. 3). We did not infer any biome shift in the remaining 11 genera, while these represent some of the most species-rich genera (e.g., *Berlinia*, *Gilbertiodendron, Anthonotha,* and *Englerodendron*).

**Figure 3.**
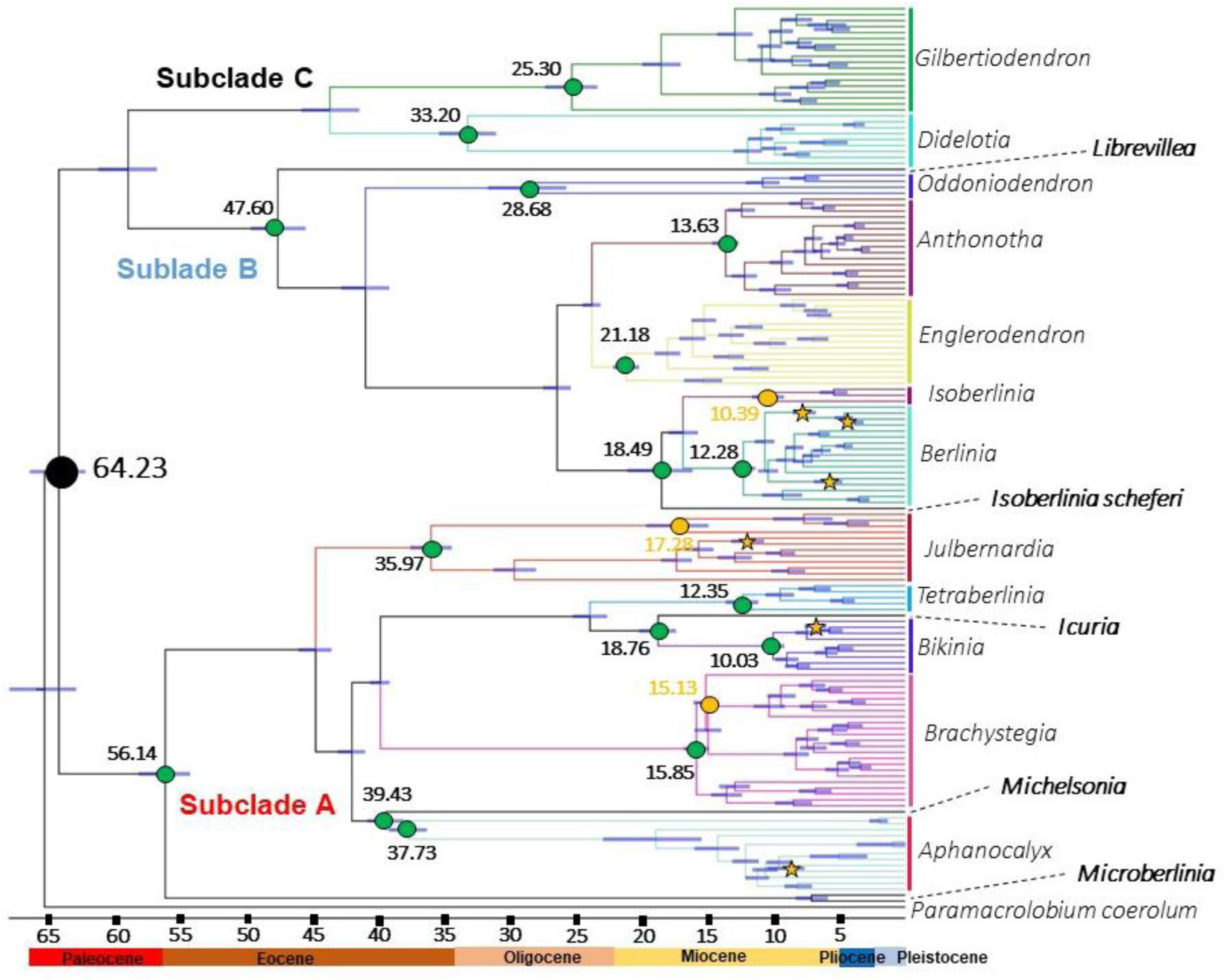
Time calibrated phylogeny of the Berlinia clade based on BEAST. The blue bars indicate 95% posterior distributions. The green circles indicate the time of the origin (crown age) of the main forest lineages (genera) and the orange circles the time of the origin of savanna lineages that diversified. Orange stars indicate shifts from forest to savanna habitats without further diversification in the savanna biome. Genera in bold indicate the paleo-endemic lineages (most monotypic).

### (c) Timing of biome shifts and its distribution among the major lineages within the Berlinia clade

Our dating analyses indicate that the Berlinia clade represents an ancient lineage that likely originated in the Paleocene (64 Ma), where 50% of its current genera originated before the Miocene (Fig. 3 and Table S4). The earliest shifts to the savanna biome occurred in the Miocene, in *Julbernardia* at about 17 Ma, in *Brachystegia* at about 15 Ma, and in *Isoberlinia* at 10 Ma. These three early biome shifts are associated with the pattern where we inferred one shift to the savanna biome with subsequent diversification within this biome. Thus, as the forest cover decreased, it seems that shifting to the newly available niche was only exploited early in the evolution of the Berlinia clade, and limited to subclades A and B. The other shifts to the savanna biome occurred in the mid-Miocene to early Pliocene in *Bikinia*, *Julbernardia*, *Aphanocalyx*, and *Berlinia*. These more recent shifts to the savanna biome are characterized by a single shift without further diversification (Fig. 3). We found phylogenetic signal for biome type with the two metrics we employed to assess non-random distribution: δ = 190.07 (p-value = 0) and D =-0.34 (p-value = 0). These results suggests that biome type is conserved in the Berlinia clade. However, our aggregation analysis suggests that the capacity to shift biomes (biome lability) is randomly distributed in the Berlinia clade at the genus level (p-value = 0.099).

### (d) Diversification rates within the forest and savanna lineages

#### (i) Time-dependent diversification

Our analyses suggest an overall increase in species number within the Berlinia clade (Fig. 4A). Of the six models tested with RPANDA, we found a constant speciation rate with no extinction as the best supported model for our data set (Table 1). The Lineages Through Time (LTT) plot indicates an increase of lineages since the early Eocene, with a marked increase from the mid-Miocene to the present (Fig. 4B). A similar trend on net diversification rates was inferred with BAMM from the Miocene (Fig. 4C), due to an increased speciation rate with a constant extinction rate (Fig. S6-S9). All our time dependent diversification analyses suggest an overall increase in net diversification within the Berlinia clade, both in forest and savanna lineages, suggesting that both groups might have exploited additional niche spaces (e.g., pollinators, soil types) within the forest and savanna biomes.

**Figure 4.**
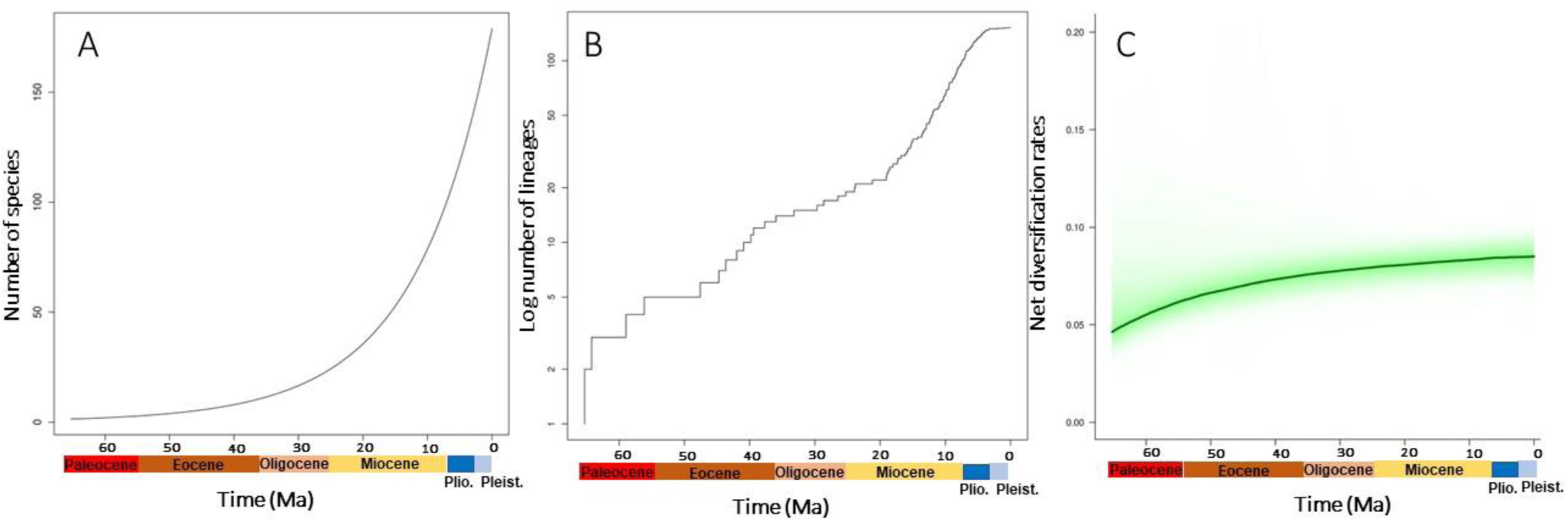
Diversification rates of the Berlinia clade based on (A) RPANDA, (B) LTT plots and (C) BAMM

**Table 1.** Results of the models tested with RPANDA in the Berlinia clade. Columns in bold indicate the best-fitted model for the data. LH, log-likelihood; AICc, corrected Akaike Information Criterion; ΔAICc, change in AICc compared with the model with the lowest AICc; Models’ parameters: λcon, speciation constant; λexp, speciation exponential; λlin, speciation linear; μ0, no extinction; μcon, constant extinction.

| Models | LH | AICc | ΔAICc |
| --- | --- | --- | --- |
| **Time-dependent** |  |  |  |
| $\lambda_{con}$ and $\mu_0$ | **-326.45** | **654.45** | 0.00 |
| $\lambda_{exp}$ and $\mu_0$ | -326.18 | 656.45 | 2.00 |
| $\lambda_{lin}$ and $\mu_0$ | -326.12 | 656.33 | 1.88 |
| $\lambda_{con}$ and $\mu_{con}$ | -326.54 | 655.11 | 0.66 |
| $\lambda_{exp}$ and $\mu_{con}$ | -326.18 | 658.53 | 4.08 |
| $\lambda_{lin}$ and $\mu_{con}$ | -326.25 | 656.58 | 2.13 |

#### (ii) Biome and trait-dependent diversification rates

To test the effect of the two biomes (forest and savanna) and the lability to shift these biomes at the genus level (labile and not labile) on diversification rates, we performed the binary state speciation and extinction (BiSSE) approach using both a ML and a MCMC method. We did not find significant differences between a constant rate and an association with the biome (α = 0.05, p = 0.305) or the lability to shift between biomes (α = 0.05, p = 0.172), suggesting that these two attributes do not seem to be associated with changes in diversification rates. The posterior probabilities of the MCMC of speciation and extinction rates for both traits overlap and indicate no difference with a constant rate of speciation in this clade (Fig. S10).

## 4. Discussion

### (a) Phylogenetic niche conservatism, but random distribution of biome niche lability

Phylogenetic niche conservatism (i.e., the retention of ancestral ecological traits and environmental distribution following speciation) is considered a major determinant of global biodiversity patterns. Evidence at the community level [67,68] as well as large geographic scale analyses suggest a high prevalence of niche conservatism [1,2,4]. We found a similar trend within the Berlinia clade, as most of the genera within this group (11 out of 16) have diversified within the same biome. In contrast to previous analyses based on genus-level phylogeny [12], our analysis provides a more detailed picture of the distribution of biome lability of African tropical trees. Our dating and ancestral reconstruction analyses within the group suggests that the Berlinia clade likely originated in the forest biome during the Paleocene (ca. 64 Ma), with the divergence of the three main subclades (A-C) in the Eocene. All this suggests that several genera within subclades A and B had expanded former distributions, retained their ancestral biome over long evolutionary time frames, and have followed the contraction of this biome. For example, there are no extant members of *Anthonotha* and *Englerodendron* in Ethiopia or Kenya, where early Miocene fossils of these genera were found, so that these genera do not seem to have exploited the availability of new drier habitat types in this geographic region [56,57]. Forests in East Africa were progressively disrupted starting in the early Miocene and often gradually replaced by deciduous forest and woodlands [53]. All extant species of *Anthonotha* and *Englerodendron* are currently distributed in the Guineo-Congolian region in central Africa [40,41]. Our results thus suggest that there are some lineages (genera) within the Berlinia clade that seem to inherently lack the capacity to exploit drier new available habitats, however this capability seems to be randomly distributed in this group. We also found no reversals to the forest biome once a lineage has shifted to savanna. Although our biome classifications might have missed some additional biomes (e.g. montane and seasonally dry forest) that were not captured, our main conclusions will not be affected if some of the species we included as forest or savanna and later reclassified into additional biomes.

### (b) Timing of biome shifts and diversification patterns in the Berlinia clade

Previous age estimates of the Berlinia clade based on several fossil calibrations within Detarioideae range between 15.4 Ma to 48.4 Ma [20,69,70]. These previous dating analyses included a limited representation of these genera and either used a secondary calibration point or a single fossil [19]. We inferred older estimates in the Berlinia clade compared to previous studies (sometimes by a factor of three). The inclusion of several Berlinia clade fossils in the current study and a more complete representation of species increased the reliability of the age estimates of the origin of all 16 genera within this clade. The recent report of fossils unequivocally assigned to the Berlinia clade from the early Eocene further supports the age estimates in our study. For example, fossils assigned to the stem of genera in the Berlinia subclade B (*Striatopollis catatumbus*) have been reported from the early Eocene (Ypresian) 51.9 Ma [50], as well as fossils from Berlinia clade A (*Peregrinipollis nigericus*) [51]. Also, well-preserved leaf/lamina fossils have been recently reported from Ethiopia of prehistoric species of forest lineages (*Anthonotha shimaglae* and *Englerodendron mulugetanum*) from the early Miocene (21.7 Ma) [56,57]. Our dating analyses question the hypothesis that the diversification of legumes into six subfamilies started after the Cretaceous-Paleogene boundary mass extinction (66 Ma) [71], although it remains compatible with some of the clock models explored by [71], which estimated the crown age of legumes at 75-80 Ma. Hereafter, we interpret our results assuming that fossil calibrations in the Berlinia clade are more reliable than previous ones, although the timing of diversification of legumes and their subfamilies remains an open question.

Our findings suggest nine independent shifts from forest to savanna, following two different patterns of biome evolution, with the earliest shifts occurring during the Miocene and followed by diversification in both savanna and forest lineages. However, we did not find evidence that biome shifts have promoted higher diversification rates in the Berlinia clade, similar to other findings of habitat shifts in Melastomataceae [72] and Malvaceae [73]. It thus seems that both forest and savanna lineages in the Berlinia clade have been able to exploit additional characteristics of the main biomes that we were not able to capture with our broad classification.

### (c) Monotypic forest paleoendemic genera with lower net diversification rates

Our analyses revealed four monotypic genera, as well as *Microberlinia* with two extant species that originated before the mid-Miocene (Fig. 3). These five genera are all restricted to the forest biome and most of them with limited distribution in tropical Africa (e.g., *Librevillea klanei* is restricted to West tropical Africa, *Isoberlinia scheffleri* is endemic to Tanzania, and *Icuria dunensis* is restricted to Mozambique). Our BAMM analyses do not indicate that these lineages have experienced higher extinction rates than other lineages in the Berlinia clade. Overall, our results suggest that the Berlinia clade has experienced low extinction rates, including the paleoendemic lineages (mostly monotypic genera) we identified (Figs. S7-S9). However, these forest paleoendemic lineages seem to have lower net diversification rates than lineages that shifted to new habitats (Fig. S9). *Paramacrolobium coeruleum*, sister to the Berlinia clade, is also a monotypic genus native to the forest biome but with a much wider distribution than *Librevillea, Icuria, Microberlinia, Michelsonia,* and *Isoberlinia scheffleri*. The current distribution of *P. coeruleum* includes western, central, and eastern Africa, from Guinea in the west to Kenya and Tanzania in the east, and south to Angola [74]. It is thus plausible that the ancestral lineage of the Berlinia clade and *P. coeruleum* also inhabited the forest biome. We found that the diversification rates were lower before the early Miocene in the entire Berlinia clade (Fig. 4b), and that forest lineages (e.g., *Berlinia* and *Anthonotha*) and savanna lineages (e.g., *Brachystegia*) that originated after this period seem to have increased net diversification rates. Thus, our results support previous reports showing that the lowland rain forest region in tropical Africa is characterized by old lineages that have persisted in these stable environments [15].

Our analyses of this endemic African tropical tree lineage suggest an uneven distribution in the capacity of these genera to exploit the availability of biomes. Despite the long evolutionary history of the Berlinia clade and the availability of new habitats during the multiple instances of forest contraction and expansion since the Miocene [13], some genera have only persisted in the forest biome. This heterogeneity is also evident in the location in the phylogeny of species-rich lineages that exploited both biomes identified in this study (forest and savanna), as the overall extension of the forest contracted since the mid-Miocene [10]. Thus, our study identified lineages that are potentially more susceptible to reductions or modifications of the forest biome due to future climate change and land use alterations. For example, future land use and landcover (LULC) projections under the RCP 8.5 (“worst case scenario”) predicts a 5.6% area reduction of the Congolian rainforest in Central Africa due to cropland replacement, and an 8% due to climate change [7]. These fast modifications of the forest biome are more likely to affect these paleo-endemic taxa. Under these predictions, these forest paleoendemic genera might be more susceptible to habitat modifications. Additional fine-scale analyses at the population level within each species will provide a more detailed picture of the genetic diversity, demographic history, and evolutionary potential for these paleo-endemic taxa.

## Ethics

This work did not require ethical approval from a human subject or animal welfare committee.

## Data accessibility

SRA sequences are deposited in the NCBI Bioproject PRJNA472454.

## Supplementary material is available online

Bait sequences and alignments have been deposited in Dryad Data Repository at XXX.

## Declaration of AI use

We have not used AI-assisted technologies in creating this article.

## Author Contributions

D.I.O.: Conceptualization; Data curation; Formal analysis; Writing – original draft; Writing – review & editing. A.P.G.: Formal analysis; Visualization; Writing – review & editing. G.D.: Formal analysis; Visualization; Writing – review & editing. S.C.: Conceptualization; Data curation; Formal analysis; Writing – review & editing. A.D.P.: Investigation; Methodology; Writing – review & editing. T.M.: Methodology; Writing – review & editing. S.V.A.: Resources; Writing – review & editing. A.B.: Resources; Writing – review & editing. F.F.: Funding acquisition; Supervision; M.E.: Resources; Writing – review & editing. O.H.: Conceptualization; Funding acquisition; Investigation; Project administration; Resources; Writing – review & editing.

## Acknowledgments

We thank the staff at the Meise and Kew herbaria for their support during the visits and collection of material.

## Supporting information

**Figures**

**Figure S1.** Phylogenetic relationships in the Berlinia clade using RAxML using the individual genes separately. Values next to branches indicate 500 bootstrap values.

**Figure S2.** Phylogenetic relationships in the Berlinia clade using RAxML using the concatenated matrix of all genes. Values next to branches indicate 500 bootstrap values.

**Figure S3.** Phylogenetic relationships in the Berlinia clade using IQ-TREE using the concatenated matrix of all genes. Values next to branches indicate 1000 fast bootstrap values.

**Figure S4.** Phylogenetic relationships in the Berlinia clade using ASTRAL. Values next to branches represent posterior probabilities.

**Figure S5.** Phylogenetic relationships in the Berlinia clade using Mr. Bayes using the concatenated matrix of all genes. Values next to branches indicate Bayesian values.

**Figure S6.** Extinction, speciation, and net diversification rates of the Berlinia clade as inferred using BAMM.

**Figure S7.** Extinction rates of the Berlinia clade as inferred using BAMM.

**Figure S8.** Speciation rates of the Berlinia clade as inferred using BAMM.

**Figure S9.** Net diversification rates of the Berlinia clade as inferred using BAMM.

**Figure S10**. BiSSE analyses of the Berlinia clade.

**Tables**

**Table S1**. List of specimens used in this study.

**Table S2**. Statistics of the target regions captured in the Berlinia clade using the Detaroideae bait.

**Table S3**. Biome classifications of the Berlinia clade.

**Table S4**. Age estimates of the genera within the Berlinia clade.

**Table S5**. Fossils used in the calibration of the phylogenentic tree in the Berlinia clade.

**Table S6**. Strategy used to estimate the lambda distribution values for the dating analysis in the Berlinia clade.

